# Protein phosphatase SppA regulates apical growth and dephosphorylates cell polarity determinant DivIVA in *Streptomyces coelicolor*

**DOI:** 10.1101/2021.03.01.433328

**Authors:** Fanny M. Passot, Stuart Cantlay, Klas Flärdh

## Abstract

Bacteria that exhibit polar growth, i.e. build their peptidoglycan cell walls in restricted zones at cell poles, often show large morphological diversity and plasticity. However, their mechanisms for regulation of cell shape and cell wall assembly are poorly understood. The Gram-positive *Streptomyces* bacteria, like other Actinobacteria, depend on the essential coiled coil protein DivIVA for establishment of cell polarity and direction of polar growth. Streptomycetes grow as filamentous hyphae that exhibit tip extension. New hyphal tips are generated by lateral branching. Cell shape is largely determined by the control of cell wall growth at these hyphal tips. The Ser/Thr protein kinase AfsK is involved in controlling polar growth and directly phosphorylates DivIVA. Here, we identify a protein phosphatase in *Streptomyces coelicolor*, SppA, that dephosphorylates DivIVA *in vivo* and *in vitro* and affects growth and cell shape. An *sppA* mutant shows reduced rate of hyphal tip extension, altered hyphal branching patterns, and exhibits frequent spontaneous hyphal growth arrests, all contributing to the unusually dense mycelial structure and slow growth rate that characterize *sppA* mutants. These phenotypes are largely suppressed in an *afsK sppA* double mutant, showing that AfsK and SppA partially affect the same regulatory pathway and share target proteins that are involved control of polar growth in *S. coelicolor*. Strains with a non-phosphorylatable mutant DivIVA were constructed and confirm that the effect of *afsK* on hyphal branching during normal growth is mediated by DivIVA phosphorylation. However, the phenotypic effects of *sppA* deletion are independent of DivIVA phosphorylation and must be mediated via other substrates. Altogether, this study identifies a PPP-family protein phosphatase directly involved in the control of polar growth and cell shape determination in *S. coelicolor* and underscore the importance of eukaryotic-type Ser/Thr phosphorylation in regulation of growth and cell envelope biogenesis in Actinobacteria.

**Author summary:** Gram-positive bacteria often use eukaryotic-like Ser/Thr protein phosphorylation in the regulation of central processes related to growth and cell cycle progression. Members of the Actinobacteria, including pathogenic mycobacteria and antibiotic-producing streptomycetes, exhibit a distinctive mode of polar growth, with cell wall synthesis being restricted to zones at cell poles and directed by the essential cell polarity determinant DivIVA. Previously, we have shown that *Streptomyces coelicolor* modulates its machinery for polar growth and cell shape determination via the Ser/Thr kinase AfsK, which phosphorylates DivIVA. Here, we identify a phosphoprotein phosphatase that targets DivIVA and reverses the AfsK-mediated phosphorylation. We show that this phosphatase has strong effects on polar growth and cell shape, including hyphal branching and the rate of tip extension. The effects are to a large extent dependent on the kinase AfsK but do not require DivIVA phosphorylation as such. The results identify a new regulator of polar growth in streptomycetes, and highlight the importance of Ser/Thr phosphorylation in regulation of actinobacterial growth and cell shape determination.

## Introduction

Bacteria are constantly adjusting their growth and cell replication in response to environmental conditions. The ability to regulate patterns of cell wall synthesis, cell division, cell shape determination, and cell envelope properties in response to external and internal signals is likely crucial for the competitive success of bacteria in their natural environment, and also affects important features like virulence and sensitivity to antibiotics [1–3]. The pathogen *Mycobacterium tuberculosis* provides good examples of these types of regulation in its adaptation of cell envelope biogenesis, both during the progression of infection in the host and during growth and stationary phase under laboratory conditions [4]. A related group of bacteria, *Streptomyces* spp., exhibit a significant morphological plasticity and adjust their cell shape in accordance with growth conditions. The streptomycetes are antibiotic-producing soil bacteria that like the mycobacteria belong to the Gram-positive phylum Actinobacteria [5]. Their morphological plasticity is intimately connected to their highly polarised mode of growth. In similarity to other members of the Actinobacteria, streptomycetes assemble their peptidoglycan cell walls in restricted zones at the cell poles [6–8]. This polar growth is directed in a fundamentally different way from how other rod-shaped bacteria, like *Escherichia coli*, *Bacillus subtilis*, and *Caulobacter crescentus*, control extension of their peptidoglycan (PG) sacculi. In these latter organisms, short filaments of actin-like MreB proteins organise the enzyme complexes that assemble new peptidoglycan into the lateral walls of the cells [9, 10]. Polarly growing Actinobacteria do not employ MreB as scaffold for PG synthesis enzymes. Instead, they rely on a protein complex formed by the coiled-coil protein DivIVA for orchestrating PG synthesis exclusively at the growing cell poles [7, 11–14]. In the pleiomorphic rod-shaped cells of mycobacteria and corynebacteria, the DivIVA-dependent PG synthesis that lead to cell elongation is established at the cell poles that are generated by cell division [15]. In contrast, polar growth in streptomycetes is uncoupled from cell division, and the cell poles where growth is established are instead generated *de novo* by lateral branching, leading to growth as branching and infrequently septated hyphae that grow by *bona fide* tip extension, very similar to the mode of growth of filamentous fungi [16, 17].

The shape of *Streptomyces* hyphae, including growth direction, curvature, and width of the cells, is laid down during cell wall synthesis at the hyphal tips, and is strongly affected by the apical DivIVA-containing protein complexes, which are referred to as polarisomes [7, 11, 18–20]. Further, the frequency and positioning of hyphal branches are mechanistically determined through a process of polarisome splitting and deposition of daughter polarisomes along the lateral walls of extending hyphae [21]. Thus, the regulatory mechanisms by which streptomycetes adjust their hyphal growth and mycelial morphology are likely acting on the polarisome and the activities at the hyphal tips but remain poorly understood. However, one pathway for regulation of hyphal growth and morphology has been identified in *Streptomyces coelicolor* that involves phosphorylation of the polarisome protein DivIVA by the Ser/Thr protein kinase AfsK [22, 23].

Hanks-type Ser/Thr protein kinases (STPKs) are common regulators in eukaryotes, but they are also widespread and have important functions in bacteria. Particularly among Gram-positive bacteria, STPKs are involved in the regulation of growth, cell cycle-related processes, and cell envelope biogenesis [24–29]. Most Gram-positives encode a transmembrane STPK with extracytoplasmic sensory domain containing PASTA (peptidoglycan and Ser/Thr kinase associated) repeats that controls processes related to cell wall growth, cell division and cell shape. PASTA domains interact with the peptidoglycan precursor lipid II or with peptidoglycan fragments [30–32]. Among members of phylum Firmicutes, the kinase StkP in *Streptococcus pneumonia*e targets a range of proteins affecting growth, cell wall homeostasis, and cell shape, including key cell shape and division proteins DivIVA and MapZ [33–37]. A similar kinase affects cell shape and division in *Streptococcus pyogenes* and *Streptococcus suis* [38, 39]. In *Staphylococcus aureus*, the PASTA-domain STKP, referred to as Stk or PknB modulates different aspects of cell envelope biogenesis [40, 41]. The *Bacillus subtilis* kinase PrkC has several functions, and among its targets are proteins related to control of cell wall metabolism [42, 43], and also the kinase PrkA in *Listeria monocytogenes* affects cell wall homeostasis [44].

Actinobacteria encode large numbers of STPKs in their genomes, with *Mycobacterium tuberculosis* having 11 STPKs, *Corynebacterium glutamicum* having four, and *Streptomyces coelicolor* encoding 34 STPKs [45–48]. Several aspects of cell envelope biogenesis in mycobacteria are directly controlled by STPKs [4, 28]. The PASTA-domain kinase PknB is essential for viability and together with the similar kinase PknA controls many processes, including cell wall synthesis and cell growth. Among the direct targets of mycobacterial PknB are the DivIVA orthologue Wag31, lipid II flippase MviN, and other cell wall-related proteins [4, 29, 49–52]. In *C. glutamicum*, STPKs phosphorylate, for example, MurC [53], coiled-coil protein RsmP [54], and FtsZ [47].

In *S. coelicolor*, DivIVA is not phosphorylated by any of the three PASTA domain kinases present in this organism [22]. Instead, the DivIVA kinase AfsK is of another type and has a C-terminal part with a putative sensory portion carrying PQQ domain repeats predicted to form a **β-**propeller structure [46]. Further, AfsK has been reported to be cytoplasmic, loosely membrane-associated [55], and to accumulate at hyphal tips [22]. The AfsK-mediated phosphorylation of DivIVA is low during growth of the vegetative mycelium, and it is strongly upregulated under conditions when peptidoglycan synthesis is blocked by bacitracin or vancomycin [22], although the stimulus or ligand sensed by AfsK is unknown. AfsK affects hyphal growth and branching both during conditions when there is a low basal level of DivIVA phosphorylation, and when AfsK activity is strongly increased [22]. Thus, it is clear that AfsK-mediated Ser/Thr phosphorylation is involved in the regulation of polar growth and cell shape determination in response to external cues in *S. coelicolor*. In order for this regulation to be reversible, a Ser/Thr protein phosphatase is needed to dephosphorylate DivIVA. The PASTA-domain kinases in Firmicutes are often associated with a protein phosphatase of the metal-dependent protein phosphatase (PPM) family, e.g. PhpP in *S. pneumoniae* or Stp in *S. aureus* [40, 56]. *M. tuberculosis* has a single Ser/Thr protein phosphatase, PstP, which also belongs to PPM family [57].

In this paper, we identify a DivIVA phosphatase in *S. coelicolor*. We demonstrate that the protein phosphatase SppA dephosphorylates DivIVA both *in vivo* and *in vitro*. In contrast to the phosphatases associated with PASTA domain STPKs in other Gram-positives, SppA belongs to the phosphoprotein phosphatase (PPP) family. It has previously been reported that SppA is a Ser/Thr/Tyr protein phosphatase, and that the *sppA* mutant grows slowly compared to the wild-type parent in *S. coelicolor* [58]. We show here that SppA indeed is involved in control of polar growth and cell shape determination in *S. coelicolor* and affects the regulatory pathway that also involves the kinase AfsK. The results underscore the importance of eukaryotic-type Ser/Thr protein phosphorylation in the regulation of growth and cell shape determination in Actinobacteria, and identifies a new type of protein involved in this regulation. In addition, as part of this study, we optimized conditions for growth of *S. coelicolor* in two dimensions, and developed image analysis methods that allow reliable quantification of the growth and morphology of the mycelium, which could be readily applied to the description of other mycelia or branching structures.

## Results

### The protein phosphatase SppA affects phosphorylation of DivIVA

The *sppA* gene (*SCO3941*) in *S. coelicolor* encodes a protein phosphatase of the PPP family with dual specificity, being able to dephosphorylate both Ser/Thr and Tyr residues [58]. It is broadly conserved in *Streptomyces* genomes. An *sppA* mutant was reported to grow slowly, suggesting that the phosphorylation status of some substrate proteins of SppA is important for vegetative growth of streptomycetes. However, no natural substrates for this phosphatase have previously been identified. Since the reduction of growth rate potentially could be related to phosphorylation of DivIVA, which is a key protein for vegetative hyphal growth in streptomycetes, we set out to test the hypothesis that SppA targets the polar growth determinant DivIVA in *S. coelicolor*. For this purpose, we constructed a strain where *sppA* is replaced by a viomycin resistance marker. In agreement with the report by Umeyama *et al.* [58], the *sppA* deletion mutant produces smaller colonies than the wild-type strain (Fig 1). This phenotype is complemented by the expression *in trans* of *sppA* (Fig 1). In contrast to what was observed by Umeyama *et al.* in a different wild-type background and on different media, we did not notice any clear difference in aerial mycelium formation or sporulation (Fig 1), reinforcing the conclusion that *sppA* mainly affects vegetative growth.

**Figure 1.**
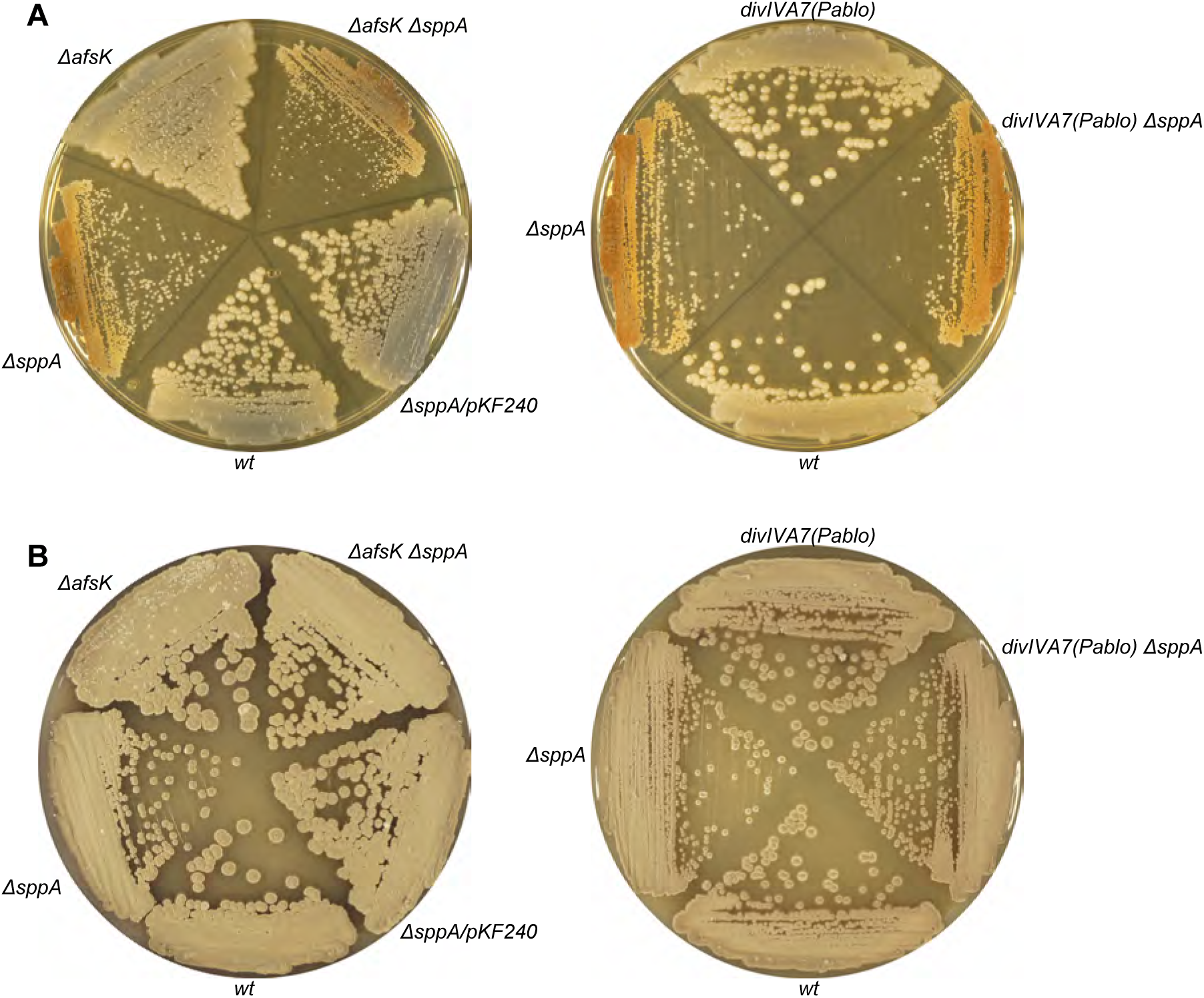
Effects of *sppA*, *afsK*, and *divIVA* mutations on colony phenotypes. **A.** Bacteria were cultivated on TSA medium for 3 days at 30°C. **B.** Bacteria were cultivated on SFM medium for 5 days (left) or 4 days (right) at 30°C. Bacterial strains and plasmids are as listed in Table 1 and Table S1.

**Table 1:**
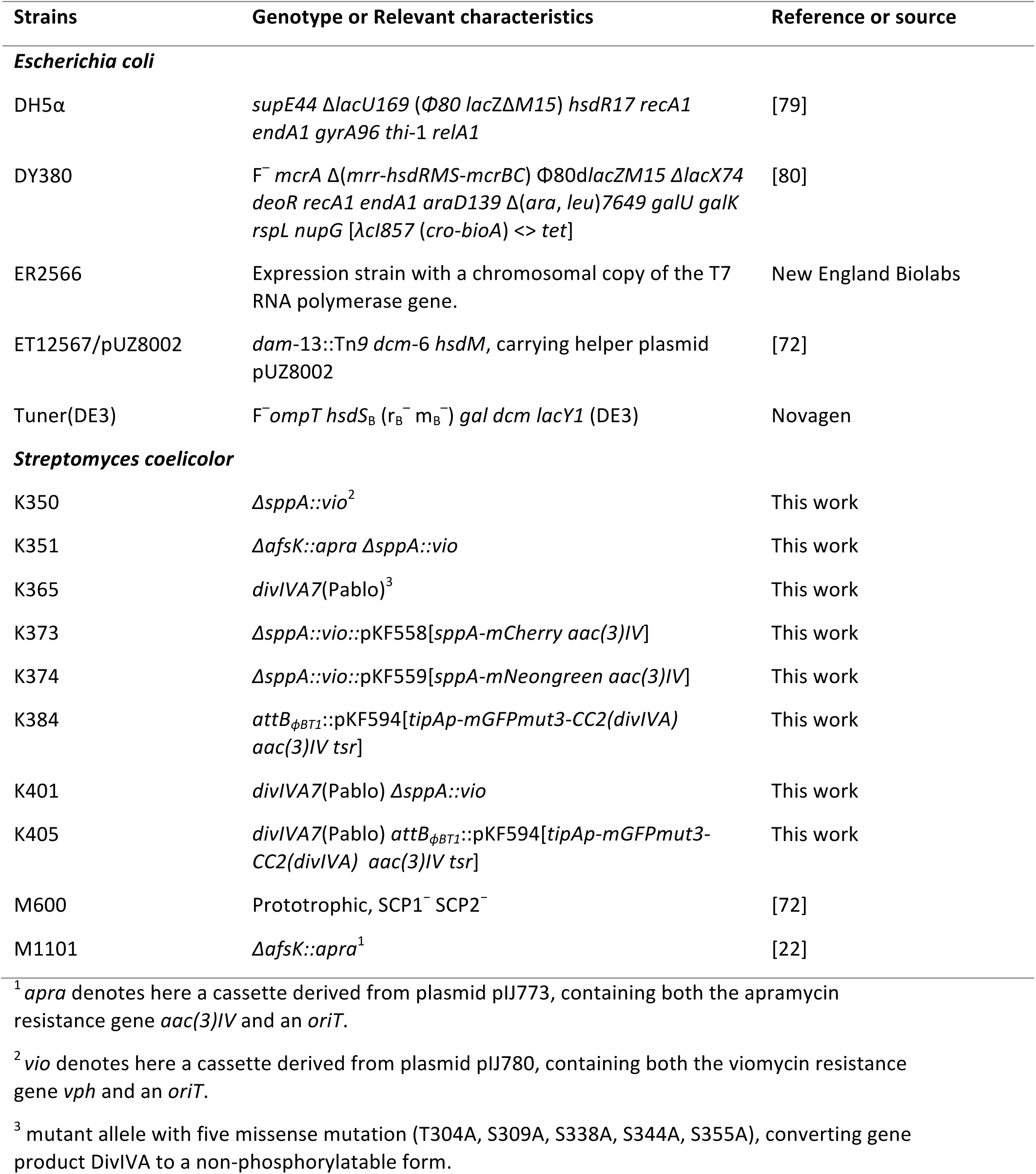
Bacterial strains used in this study.

To determine whether *sppA* affects DivIVA phosphorylation, we took advantage of the previously demonstrated electrophoretic mobility shift of the phosphorylated forms of DivIVA compared to the non-phosphorylated forms, as detectable by SDS-PAGE and Western blotting [22]. In extracts from an untreated culture of a wild type strain, DivIVA appears on the Western blot as an intense fast-migrating band, corresponding to the non-phosphorylated molecules, and a single slightly shifted weak band, corresponding to phosphorylated forms (Fig 2). Thus, in agreement with our previous report [22], DivIVA is largely non-phosphorylated under these conditions of normal growth. When peptidoglycan synthesis is blocked by bacitracin treatment for 30 min, the non-phosphorylated faster band is not visible on the Western blot and is replaced by several shifted bands, indicating that DivIVA phosphorylation increases strongly when PG synthesis is arrested (Fig. 2), also as shown previously [22]. In a growing *ΔsppA* strain, without any treatment of the culture, several shifted bands are already visible on a DivIVA Western blot (Fig 2). Upon bacitracin treatment, the fast-migrating non-phosphorylated band disappears and the shifted bands are shifted even higher, to what seems to be the most phosphorylated form (Fig 2). These data indicate that in a *ΔsppA* strain, both the fraction of DivIVA molecules that are phosphorylated and the level of phosphorylation of each molecule is increased, reaching saturation after the bacitracin treatment. We further confirmed a similar effect on DivIVA phosphorylation in a different strain background (*S. coelicolor* strain M145) and with a different type of mutation in *sppA* (a transposon insertion). This mutation gave a similar small colony phenotype and increased levels of DivIVA phosphorylation (S1 Fig, the effect is clearly seen when comparing the first wells to the left for the two strains, representing time point −10 min), as described above, and was also complemented by *sppA in trans*.

**Figure 2.**
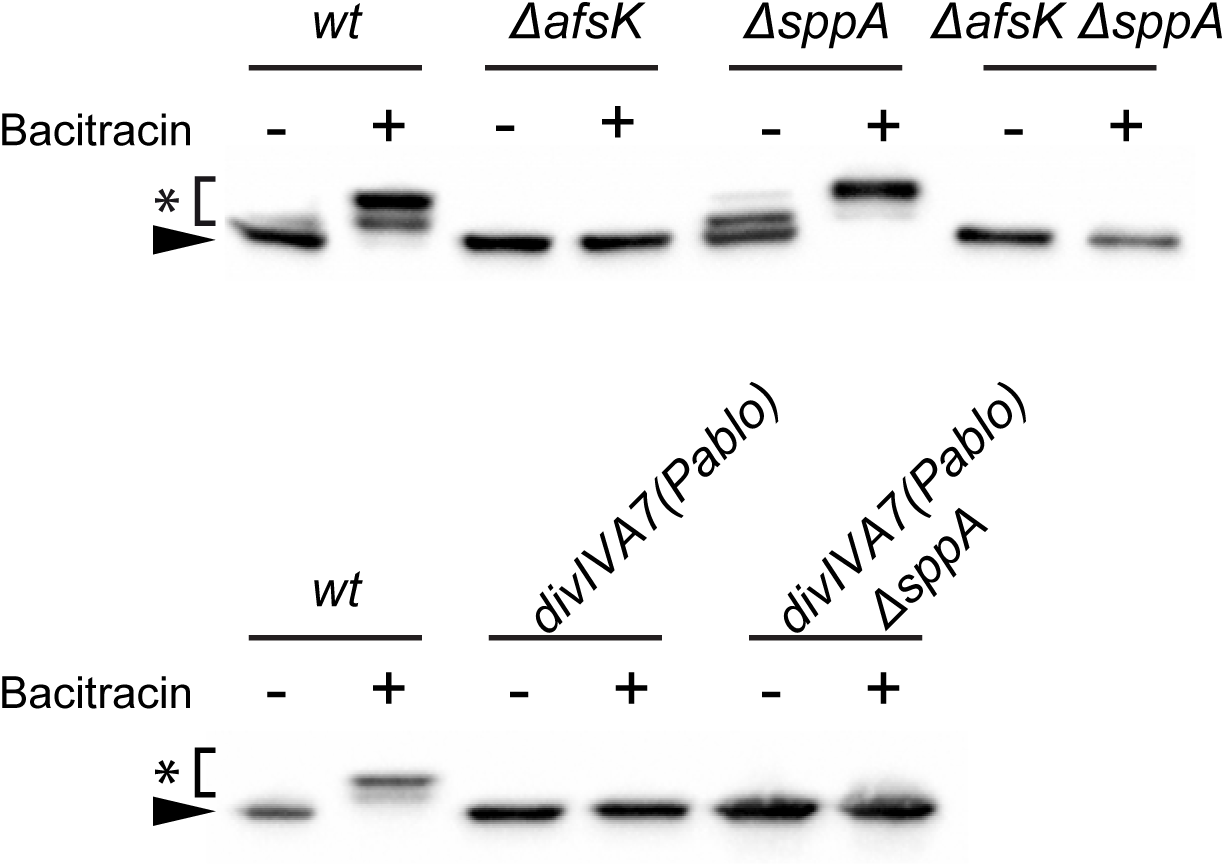
Phosphorylation of DivIVA during growth and upon arrest of peptidoglycan synthesis growth arrest. Growing cultures of *S. coelicolor* were exposed to bacitracin (50 μg/ml) for 30 min. Cells were harvested and total cell extracts were prepared, separated by SDS-PAGE and subjected to anti-DivIVA Western blots. The fast-migrating main band represents unphosphorylated DivIVA (black arrowhead), while the slower migrating bands derive from phosphorylated forms of DivIVA (marked by asterisk).

In order to monitor relative rates of dephosphorylation of DivIVA *in vivo*, we compared the persistence of the highly phosphorylated DivIVA after the cell wall synthesis inhibitor had been washed away in the wild type and *ΔsppA* strain. The phosphorylated forms of DivIVA returned to the non-phosphorylated state more slowly in the *sppA* mutant (S1 Fig), showing that the rate of dephosphorylation of DivIVA is lower in the absence of the SppA phosphatase.

### SppA can directly dephosphorylate DivIVA

Next, we asked whether purified SppA can directly dephosphorylate DivIVA *in vitro*. *S. coelicolor* SppA was purified from a recombinant *E. coli* strain, and its phosphatase activity was confirmed using colorimetric assays with p-nitrophenylphosphate (pNPP) as substrate (data not shown), in agreement with a previous report [58]. A phosphorylated version of DivIVA (phospho-DivIVA) was purified from a recombinant *E. coli* strain expressing both DivIVA and the kinase domain of AfsK, which is known to target DivIVA [22]. All detectable DivIVA in these preparations exhibited a mobility shift on SDS-PAGE gels that is typical of phosphorylated DivIVA in *S. coelicolor* (Fig 3)[22]. When phospho-DivIVA was incubated with SppA in previously established reaction conditions (see Material and Methods section), a mobility shift back to the position of non-phosphorylated DivIVA revealed protein phosphatase activity. This activity was inhibited by EDTA (Fig 3), which is consistent with the stimulation of SppA activity by divalent cations [58]. Together with the *in vivo* results described above, these data suggest that SppA indeed is a major phosphatase of DivIVA in *S. coelicolor*.

**Figure 3.**
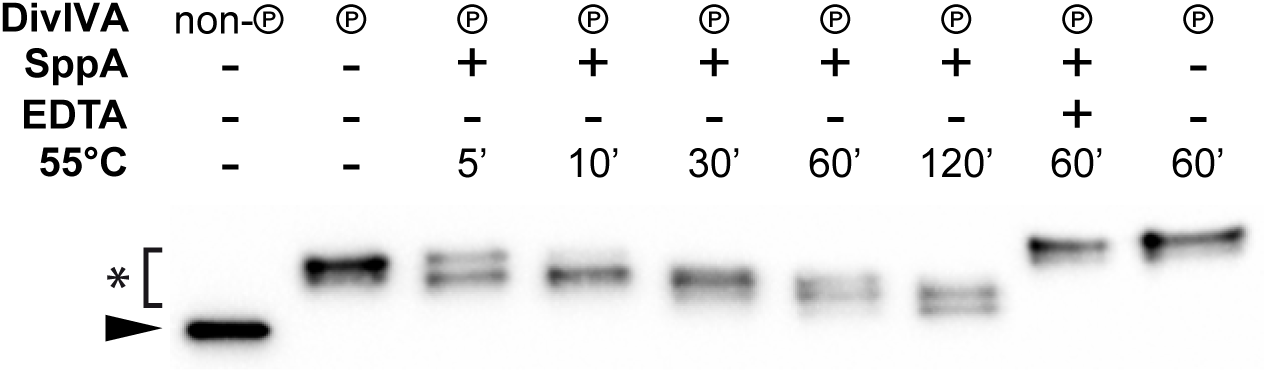
*In vitro* dephosphorylation of phospho-DivIVA by SppA. Phosphorylated DivIVA was heterologously produced in *E. coli* by co-production with its cognate kinase AfsK. Phospho-DivIVA (℗) was mixed with purified SppA in Mn^2+^-containing buffer and incubated at 55°C for 5, 10, 30, 60, and 120 min (lanes 3-7), before stopping reactions and analysing by SDS-PAGE and Western blotting with anti-DivIVA antiserum. An unphosphorylated preparation of DivIVA shows the position of the unphosphorylated form on the gel (lane 1), while the phosphorylated forms show distinctive mobility shifts (lane 2). No dephosphorylation was observed in the presence of EDTA (lane 8) or in the absence of SppA (lane 9). Black arrowhead, unphosphorylated DivIVA; asterisk, phosphorylated forms of DivIVA.

### SppA localizes throughout the mycelium during vegetative growth

Since both DivIVA and the DivIVA kinase AfsK are accumulated at the tips of growing hyphae [22], it was relevant to determine the subcellular localization of SppA. For this purpose, two fusions of *sppA* to genes for fluorescent proteins were constructed and expressed from the *sppA* promoter. Both fusion proteins complemented the phenotype of *ΔsppA* (restoring normal mycelial growth and morphology, see below), indicating that SppA is fully functional in these fusions. Microscopy observations showed a diffuse localization of both fluorescent hybrid proteins throughout the entire mycelium, with no tendency to accumulate at hyphal tips (Fig 4 and S2 Fig). Thus, SppA localization is only partially overlapping with DivIVA.

**Figure 4.**
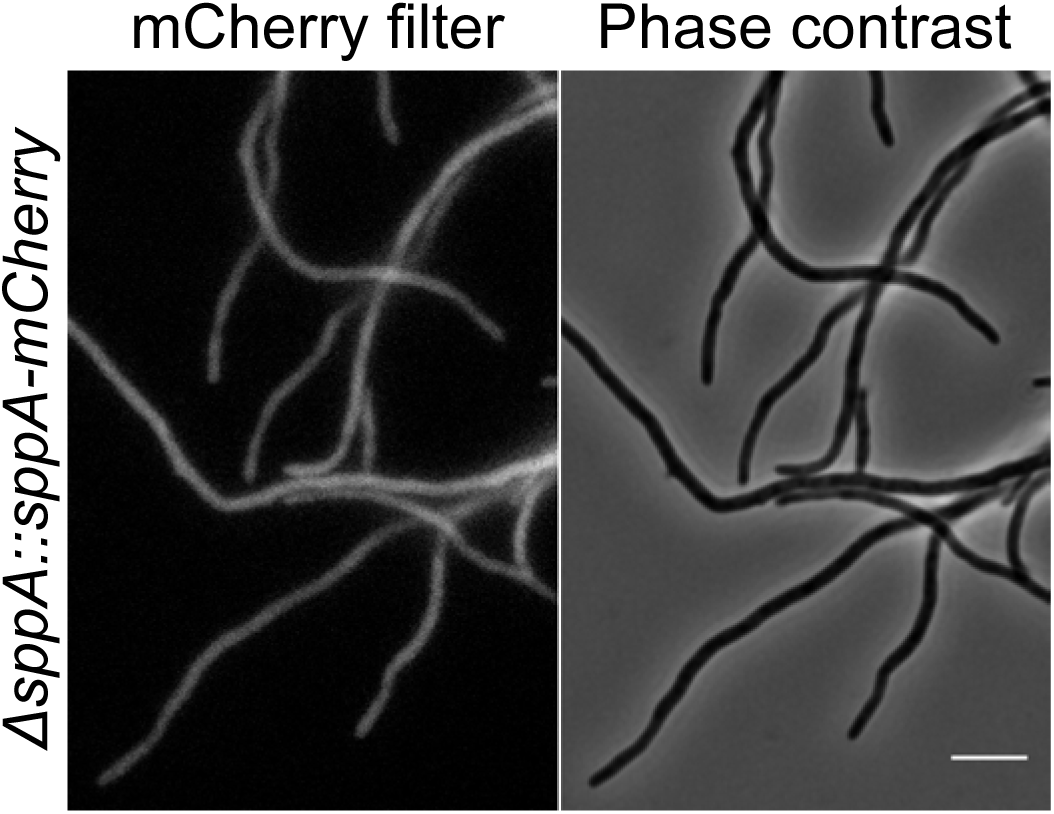
Subcellular SppA localisation. Expression of *sppA-mCherry* and *sppA-mNeongreen* fusions in strain K373. Hyphae from culture growing in TSBS medium were mounted on agarose-coated slides and fluorescence microscopy images captured with 63x objective and filter sets for detection of mCherry. Control experiments are shown in Supplementary Fig. S2. Scale bar, 5 μm.

### Deletion of *sppA* strongly affects mycelial growth and morphology

Knowing that SppA directly dephosphorylates DivIVA and affects overall rate of growth, we next tested whether SppA is involved in the control of polar growth, hyphal morphology, and hyphal branching, which are intimately connected to the DivIVA polarisome and its function at the hyphal tips. To facilitate analyses of hyphal growth patterns, mycelia were cultivated on cellophane membranes on agar plates, then transferred carefully to agarose-coated slides, and hyphal shapes were investigated by phase-contrast microscopy. Under these conditions, young mycelia from wild type cultures are spread out as loose mycelial networks with individual hyphae extending far outside the core of the mycelial clumps or pellets. In contrast, the mycelia from *ΔsppA* cultures appear denser and not as spread out, and they show only short hyphae extending outside the mycelial clumps (Fig 5). A similar phenotypic difference can be observed in liquid cultures, with the *sppA* mutant giving smaller and more dense hyphal pellets than the wild type parent. Thus, at the microscopic level, the mycelia of the *ΔsppA* strain present a clear difference in overall shape and ability to spread over a surface compared to the ones from the wild-type strain (M600).

**Figure 5.**
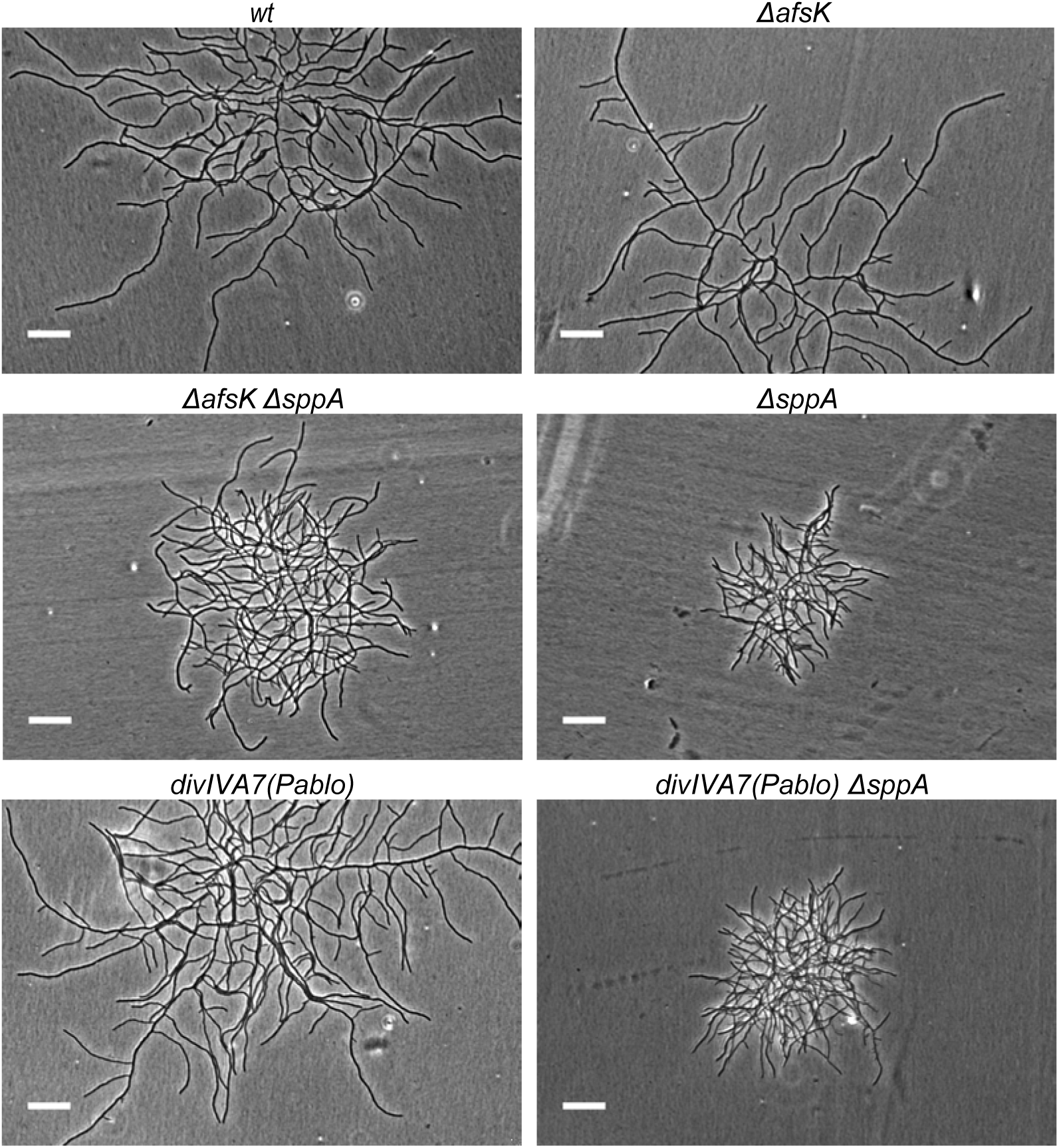
Effects on mycelial morphology. Bacteria were grown from spores on cellophane discs in a TSB-agarose chamber at 30°C. Phase contrast microscopy images of mycelia were captured 15h after inoculation. Scale bars, 20 μm. Bacterial strains are as listed in Table 1.

### Slow hyphal tip extension and frequent spontaneous growth arrests occur in *sppA* deletion mutants

To get further insight into the mechanisms causing *ΔsppA* mycelia to be denser and to form small and compact mycelial pellets, we measured several parameters of the mycelial growth and morphogenesis. In liquid cultures, as well as on agar pads, *S. coelicolor* strains form three-dimensional mycelia, which makes precise measurements difficult. To counter that, we used a microfluidic growth chamber forcing the mycelia to grow in two dimensions only (at least during the early vegetative phase), as it has previously been done for *S. venezuelae* [18, 59]. After inoculation of spores in the chamber, they are allowed to germinate in pre-germination medium for eight hours, then the medium is switched to TSB and the young mycelia are allowed to grow for six more hours before we start time-lapse imaging for up to seven hours. In this setup, the characteristic *ΔsppA* phenotype with dense mycelial clumps is observed (S3 Fig and Supplementary Movies 1 and 2), similarly to what was described above. We manually tracked the tips of the hyphae to estimate their extension rates. For the wild type strain, the central 70% of the extension rate data are distributed between 0.19 and 0.36 µm/min, whereas for *ΔsppA* these are distributed between 0.10 and 0.17 µm/min, with significantly different average values (Fig 6). During the tracking of the tips, we also noticed that hyphae stop growing much more frequently in the *ΔsppA* strain than in the wild type (Fig 7 and Supplementary Movies). 7.4% of the wild-type hyphae spontaneously stopped growing during the period of tracking, while 48.4% of the *ΔsppA* hyphae showed such spontaneous growth arrests. The slow extension of the hyphae and the high frequency of growth arrests are likely contributing to the small size of the mycelial pellets, low growth rate of cultures, and small colonies of the *ΔsppA* mutant.

**Figure 6.**
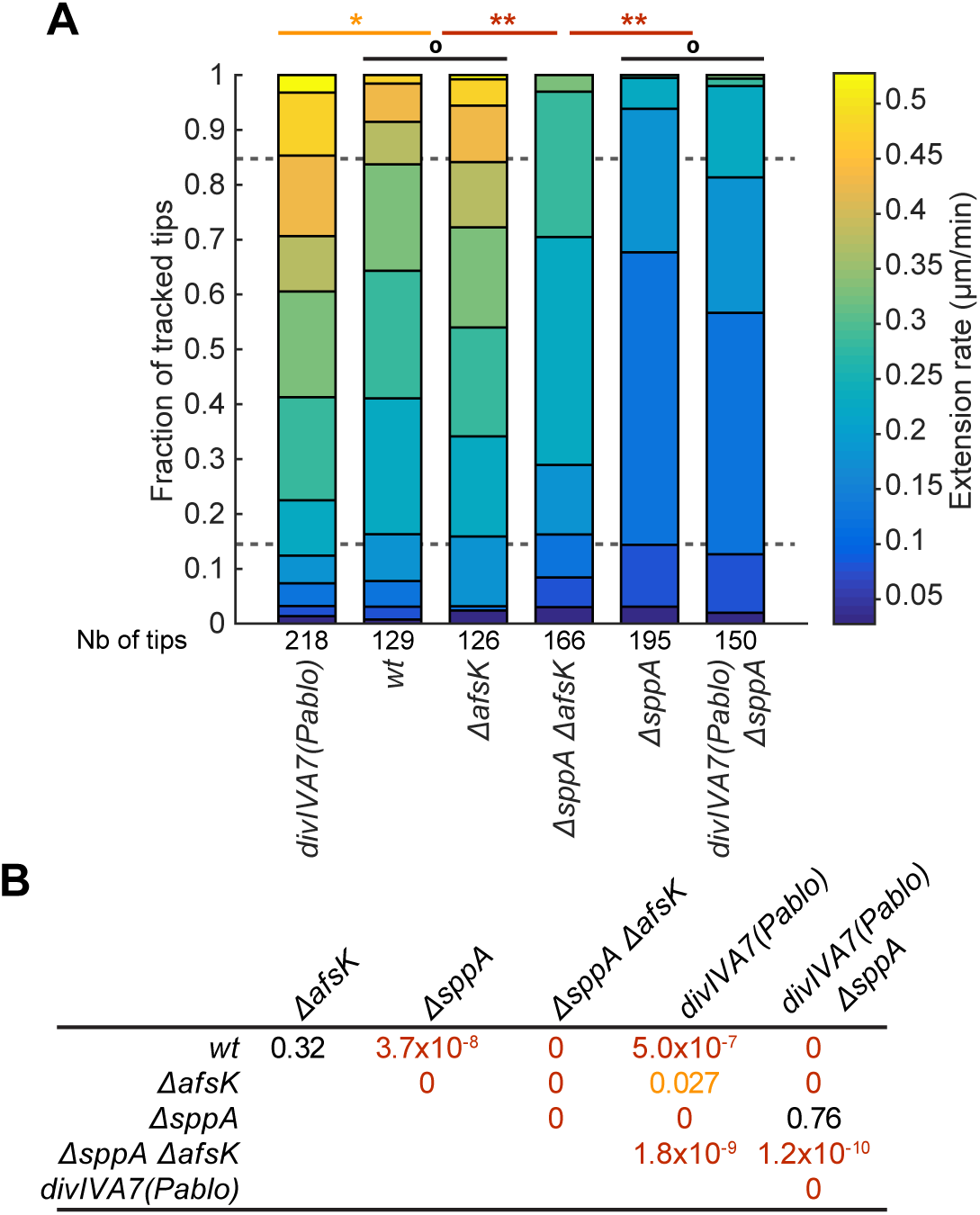
Analysis of hyphal tip extension rates. **A.** Stacked histogram of the extension rates of hyphae that were more than 1 hour old, measured on time-lapse data series from CellASIC microfluidic plate (B04A-03) in trap depth IV (0.9 μm) on growing hyphae and branches. ° non-significant difference; * significant difference with a p-value < 0.05; ** significant difference with a p-value < 10^-6^. **B.** Table of the p-values obtained through a Tukey’s pairwise test on the extension rates of the different strains. Black: non-significant difference (p-value > 0.2); orange: p-value < 0.05; red: p-value < 10^-6^.

**Figure 7.**
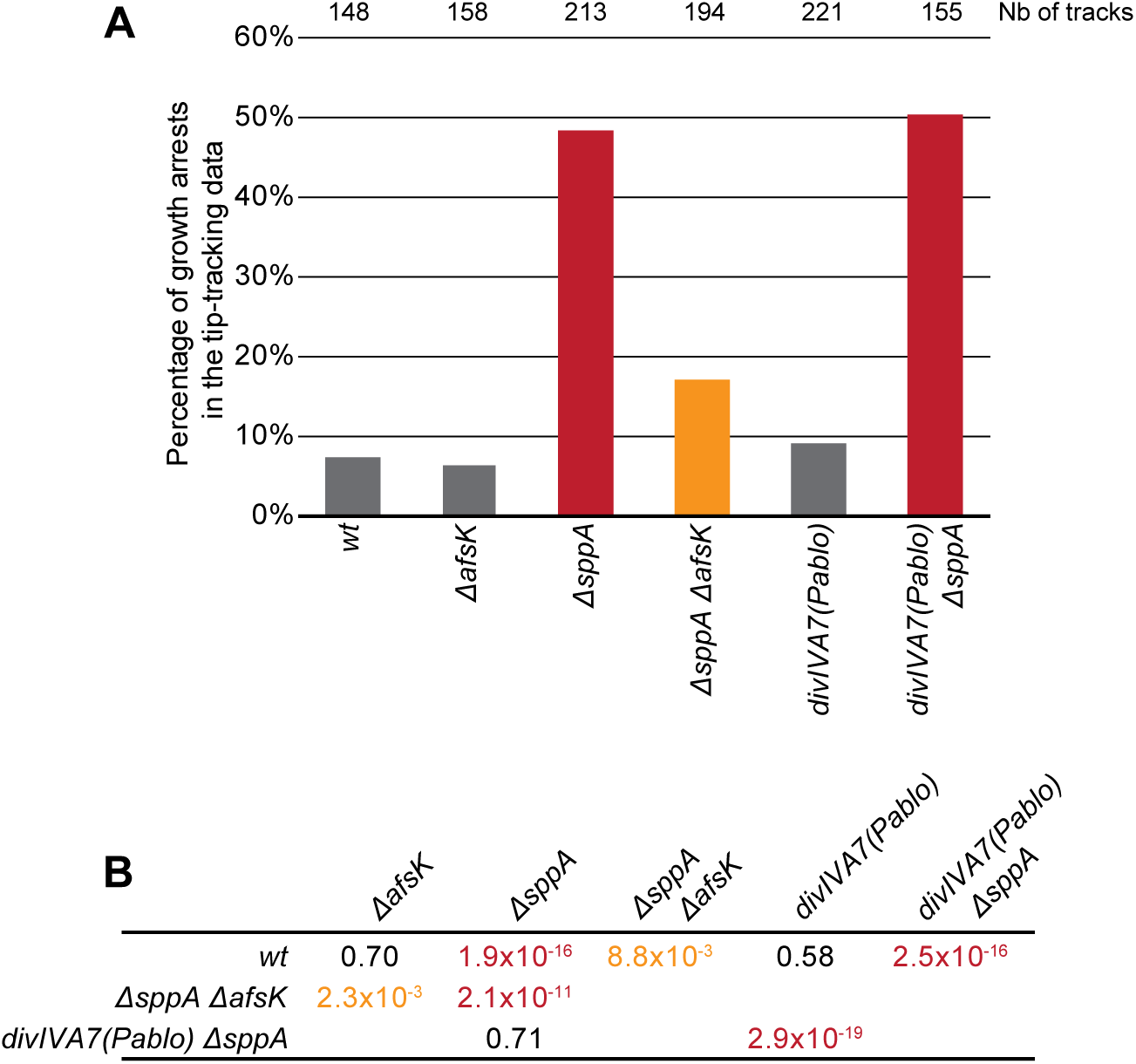
Analysis of the frequency of growth arrests of individual hyphae. **A.** Percentage of tracked hyphae whose growth stopped during the tracking in the CellASIC ONIX microfluidic system (plate B04A-03). Measurements were done on timelapse data series in trap depth IV (0.9 μm) on growing hyphae and branches. Grey bars: non-significantly different from wild type (wt); orange: significantly different from wt with a p-value < 0.05; red: significantly different from wt with a p-value < 10^-6^. **B.** Table of the p-values obtained through a Χ^2^ pairwise test on the proportion of growth arrests of the different strains. Black: non-significant difference (p-value > 0.05); orange: p-value < 0.05; red: p-value < 10^-6^.

### *sppA* deletion mutants are hyperbranching

The *ΔsppA* mycelial clumps are not only smaller but also denser than the wild type ones. This could come from a higher branching frequency (i.e. a shorter branch-to-branch distance) and/or from branching closer to the tip (i.e. a shorter tip-to-branch distance) [21]. The branch-to-branch distance was unfortunately impossible to measure due to too many crossing hyphae in the deeper parts of the mycelium and formation of new branches on old hyphae over time. On the contrary, the tip-to-branch distance can be reliably measured on the dispersed outer parts of the mycelium and can be clearly pinpointed in time. Thus, in the same time-lapse data that we used for measurements of the extension rates, we measured the distance from the tip of all new tip-proximal branches, at the time points when these branches are 1 to 4 µm long (Fig 8). These threshold sizes are needed to take into account the fact that the new branches are visible earlier if they come out of the sides of the hyphae than if they come out from the top or bottom of it. Measurements are only done on growing hyphae, as could be judged in the time-lapse series, to avoid biases due to further branching of the hyphae whose tip is not extending anymore. Under these conditions, the central 70% of the tip-to-branch distance distribution in the wild type mycelia is comprised between 12.5 and 33.5 µm, while it is comprised between 10.4 and 19.1 µm in the *ΔsppA* mycelia, with the difference being statistically significant (Fig 8). The tip-to-branch distance in the growing hyphae is thus shorter in *ΔsppA* than in the wild type. In addition, hyphae that stop elongating usually start branching abundantly (Supplementary movies, white arrows), and as growth arrests are much more frequent in *ΔsppA*, this also contributes to a higher overall frequency of branching in *ΔsppA* mycelia. In summary, the higher density of *ΔsppA* mycelial clumps (compared to the wild type) appears to be caused by a combination of lower tip extension rate, branching closer to the tip on hyphae that are growing, and more frequent growth arrests.

**Figure 8.**
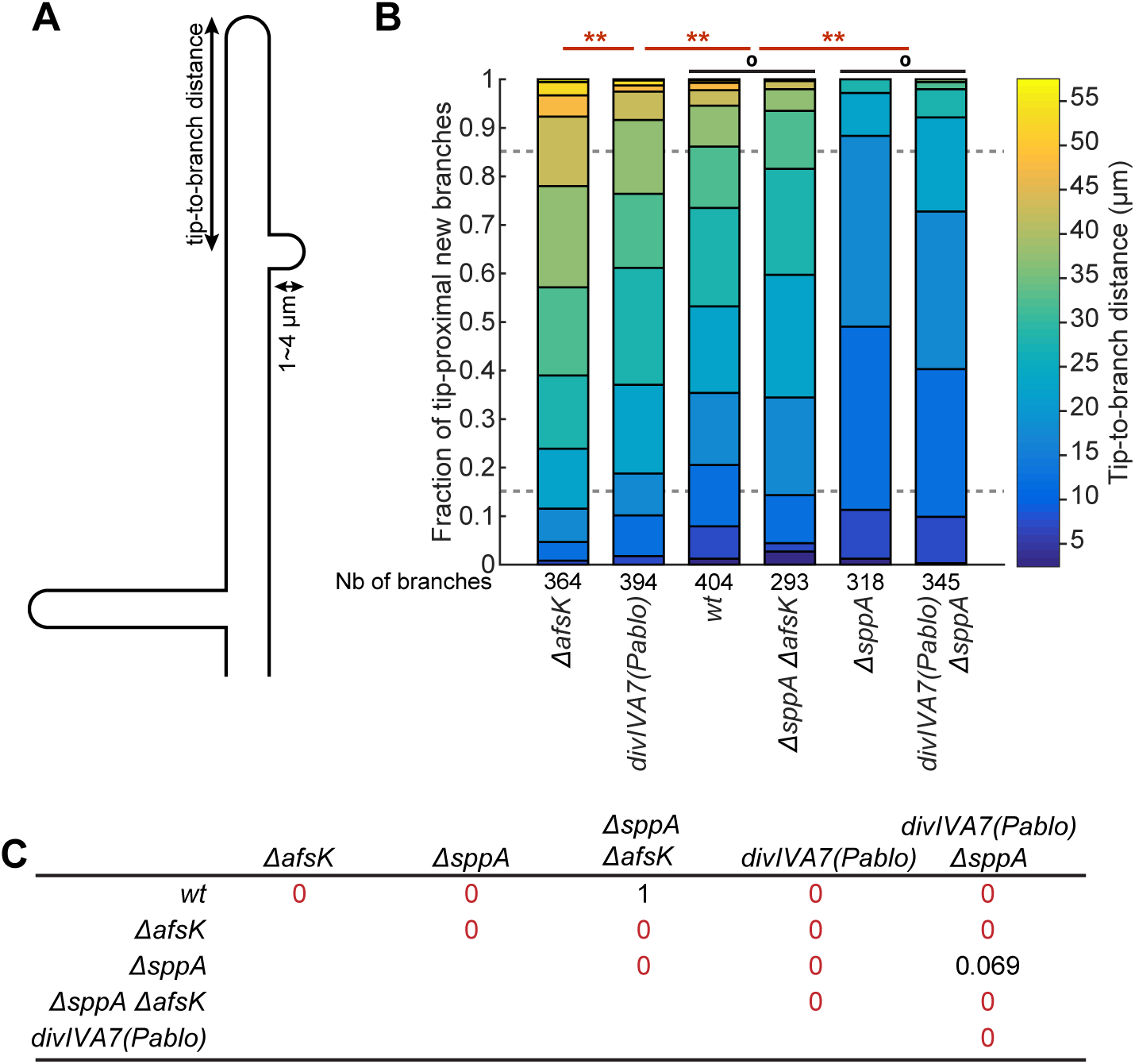
Analysis of the distribution of tip-to-branch distances. **A.** Schematics showing how the measurements were done. **B.** Stacked histogram of the distances between the tips and the tip-proximal new branches (<4 μm long) measured on timelapse data series in trap depth IV (0.9 μm) on growing hyphae and branches in the CellASIC ONIX microfluidic system (plate B04A-03). ° non-significant difference; ** p-value < 10^-6^. **C.** Table of the p-values obtained through a Tukey’s pairwise test on the tip-to-branch distances of the different strains. Black: non-significant difference (p-value > 0.05); red: p-value < 10^-6^.

### The *sppA* mutant phenotype is partially suppressed by *afsK* deletion

Since both the kinase AfsK and the phosphatase SppA target DivIVA, but affect DivIVA phosphorylation in opposite ways, we asked whether deletion of *afsK* (which abolishes DivIVA phosphorylation) could suppress any aspect of the phenotype caused by the *ΔsppA* mutation. For this purpose, the *ΔsppA::vio* allele was introduced into the previously constructed *ΔafsK* strain (M1101) [22], generating a *ΔafsK ΔsppA* double mutant (K351). We first analyzed DivIVA phosphorylation in these strains (Fig 2). As seen previously [22], both the basal level of DivIVA phosphorylation and its induction by bacitracin treatment are abolished in the *ΔafsK* strain. This is also the case in the *ΔafsK ΔsppA* double mutant, showing that *ΔafsK* is epistatic on *ΔsppA* for DivIVA phosphorylation, and further confirming that AfsK likely is the only kinase of DivIVA under the tested conditions [22].

At the colony level, while *ΔafsK* colonies are indistinguishable from the wild type, *ΔafsK ΔsppA* colonies are larger than *ΔsppA* ones on SFM medium but smaller than wild type ones (Fig 1B). This intermediate phenotype is also visible at the mycelium level (Fig 5). More precisely, while the extension rate of *ΔafsK* hyphae, similarly to the wild type, is mainly distributed between 0.20 and 0.41 µm/min (percentiles 15 and 85), the extension rate of *ΔafsK ΔsppA* hyphae is mainly distributed between 0.15 and 0.27 µm/min, significantly slower than the wild type, but also significantly faster than the *ΔsppA* hyphae (Fig 6). Similarly, while growth arrests are quite rare in *ΔafsK* hyphae (6.3%), as in the wild type (7.4%), they are more frequent in *ΔafsK ΔsppA* (17%), but not as much as in *ΔsppA* (43.4%) (Fig 7).

As shown previously, the average distance from tips to first branches in *ΔafsK* mycelia is longer than the wild type [22]. In our microfluidic setup, this is visible as tip-to-branch distances being distributed mainly between 22.1 and 42.4 µm (percentiles 15 and 85) in *ΔafsK* mycelia while they are distributed mainly between 12.5 and 33.5 µm in wild type mycelia (Fig 8). In *ΔafsK ΔsppA* mycelia, these distances are mainly distributed between 15.2 and 30.8 µm, in a distribution that is not significantly different from the wild type, once again showing an intermediate phenotype between *ΔsppA* and *ΔafsK*. Altogether, the results indicate that *ΔafsK* partially suppresses the phenotypes of *ΔsppA*, suggesting that their target proteins are at least partially overlapping.

### A mutant with a phosphoablative version of DivIVA is viable and has a similar branching pattern to the *afsK* mutant

To be able to directly test the involvement of DivIVA phosphorylation in the characteristic phenotypes of *afsK* and *sppA* mutants, we constructed a mutant *divIVA* allele that encodes a non-phosphorylatable version of DivIVA. The phosphorylated residues of DivIVA were previously identified to be T304, S309, S338, S344 and S355 [23]. By site-specific mutagenesis, the five codons corresponding to these phosphorylatable serines and threonines were replaced by alanine codons. The native *divIVA* gene in *S. coelicolor* strain M600 was then replaced by this phosphoablative mutant allele, named *divIVA7*(Pablo), through integration and excision of a counter-selectable plasmid. Depending on the site of homologous recombination for the integration of the plasmid, its excision is expected to leave the mutant allele on the chromosome in 42% of the cases and revert to the wild type in 55% of the cases, or the opposite. Out of 11 sequenced candidates after excision, 7 were carrying the mutant allele and 4 were carrying the wild type allele. The resulting *divIVA7*(Pablo) mutant strain is viable and give colonies that are indistinguishable from the wild type ones (Fig 1 and S4 Fig). On a DivIVA Western blot of cell extracts from this strain, DivIVA7(Pablo) appears as a band similar to the non-phosphorylated DivIVA, even upon bacitracin treatment (Fig 2), confirming that this mutant form of DivIVA is not phosphorylated under the tested conditions.

At the microscopic level, the mycelium morphology of the *divIVA7*(Pablo) strain is largely similar to the wild type (Fig 5). The analysis of time-lapse data of this strain indicates that growth arrest frequency (9.1%) is not significantly different from the wild type (7.4%; Fig 7). The tip extension rate of *divIVA7*(Pablo) is marginally faster than the wild type, with the central 70% of the data distributed from 0.21 to 0.45 µm/min (Fig 6). Importantly, however, the distribution of tip-to-branch distances is shifted towards longer distances in *divIVA7*(Pablo) (mainly 17.3 to 37.9 µm), mirroring the effect of *ΔafsK* on branching patterns. This finding suggests that the reported effect of *afsK* deletion on hyphal branching patterns during vegetative growth [22] is indeed mediated by the absence of DivIVA phosphorylation.

### The effects of *sppA* deletion on hyphal growth and morphology are not caused by hyperphosphorylation of DivIVA

The partial suppression of the *ΔsppA* mutant phenotype by *afsK* mutation, combined with the observation of increased level of DivIVA phosphorylation in the *ΔsppA* mutant, suggested the possibility that parts of the phenotype caused by *sppA* inactivation could be caused by DivIVA phosphorylation. If the strong phenotype of *ΔsppA* is due to a hyperphosphorylation of DivIVA, the *divIVA7*(Pablo) allele should suppress this phenotype. To test this possibility, we constructed a *divIVA7*(Pablo) *ΔsppA* strain and analysed it in the same way as described above. On a DivIVA Western blot, DivIVA7(Pablo) appears as a single fast migrating band in an untreated culture of this strain (Fig 2), indicating that DivIVA7(Pablo) is still non-phosphorylated despite the absence of SppA. Similarly, when the culture is treated with bacitracin, DivIVA7(Pablo) shows no mobility shift in the gel. Thus, the phosphoablative mutations in *divIVA7*(Pablo) prevent the hyperphosphorylation of DivIVA that is otherwise seen in *ΔsppA* mutants.

Nevertheless, the prevention of DivIVA phosphorylation does not restore normal growth to the *sppA* deletion mutant. The *divIVA7*(Pablo) *ΔsppA* colonies (Fig 1), as well as mycelial clumps formed by this strain (Fig 5), are as small as those of the *ΔsppA* mutant. Similarly, the *divIVA7*(Pablo) *ΔsppA* strain is indistinguishable from *ΔsppA* when it comes to tip extension rate (distributed mainly from 0.10 to 0.21 µm/min; Fig 6), frequency of growth arrests (50.3%; Fig 7) and tip-to-branch distances (mainly 10.8 to 22.6 µm; Fig 8). Thus, none of the growth and morphology phenotypes of *ΔsppA* is suppressed by *divIVA7*(Pablo), indicating that *sppA* affects tip growth and branching patterns by other routes than DivIVA phosphorylation, and that AfsK and SppA likely share additional substrate proteins that are involved in control of polar growth in *Streptomyces*.

### AfsK-mediated DivIVA phosphorylation is not required for the regular localization of polarisomes in growing hyphae or their disassembly upon growth arrest

Taking advantage of the *divIVA7*(Pablo) mutant, in which DivIVA is not detectably phosphorylated, we have investigated the role of AfsK-mediated phosphorylation and whether it detectably affects the DivIVA polarisomes seen at tips of growing hyphae. To get a reliable quantification of the fluorescence signal along the hyphae, we needed them to grow in two dimensions only. The CellASIC microfluidic chambers allow for that, but our previously used fluorescent fusions of DivIVA [7, 22] induced slight hyperbranching in this setup. To prevent side effects of the fluorescence tagging of DivIVA, we fused only its second coiled-coil domain (CC2), implied in oligomerisation [60], with mGFPmut3, separated by half of the long and presumably non-structured PQG domain upstream of CC2. We placed this GFP-CC2 fusion under the control of the thiostrepton-inducible promoter *tipAp*. The production of the GFP-CC2 fusion could then be induced in order to decorate the existing DivIVA complexes in the mycelia. When induced by 1 µg/ml of thiostrepton in a wild type background, the fluorescent fusion localizes similarly to what has previously been observed for DivIVA [7, 11], without affecting the branching pattern. As the CC2 domain included in this construct does not contain any of the phosphorylated sites of DivIVA, we could use the same GFP-CC2 to decorate the DivIVA7(Pablo) complexes as well as the wild type DivIVA ones. For both strains, the spores were inoculated into the growth chamber and allowed to germinate and grow for 11h before induction of GFP-CC2 production with thiostrepton. The mycelia were incubated 3h further to allow for proper expression of GFP-CC2 before imaging. We then extracted the fluorescence profile along approximately 40 hyphae starting from the tip. A control strain was also used that carries the same inducible promoter *tipAp* but without the GFP-CC2 construct. In that strain, a small focus of fluorescence is visible at the tip of the hyphae with the filter set used for GFP (Fig 9) and correspond to autofluorescence from an unknown natural product of *S. coelicolor*. In a wild type background, GFP-CC2 fluorescent signal is present mainly as a focus at the tip of the hyphae, shown on the fluorescence intensity graph as a sharp peak close to the origin (Fig 9). The same pattern is visible in a *divIVA7*(Pablo) background with no significant difference of intensity with the wild type, be it in the tip-localized focus or in the diffuse fluorescence further back in the hyphae (Fig 9). Thus, even though the *divIVA7*(Pablo) allele gives a similar effect as deletion of *afsK* on the branching pattern (longer tip-branch distance), the overall subcellular distribution of DivIVA was not detectably affected by the phospho-ablative mutations during vegetative growth.

**Figure 9.**
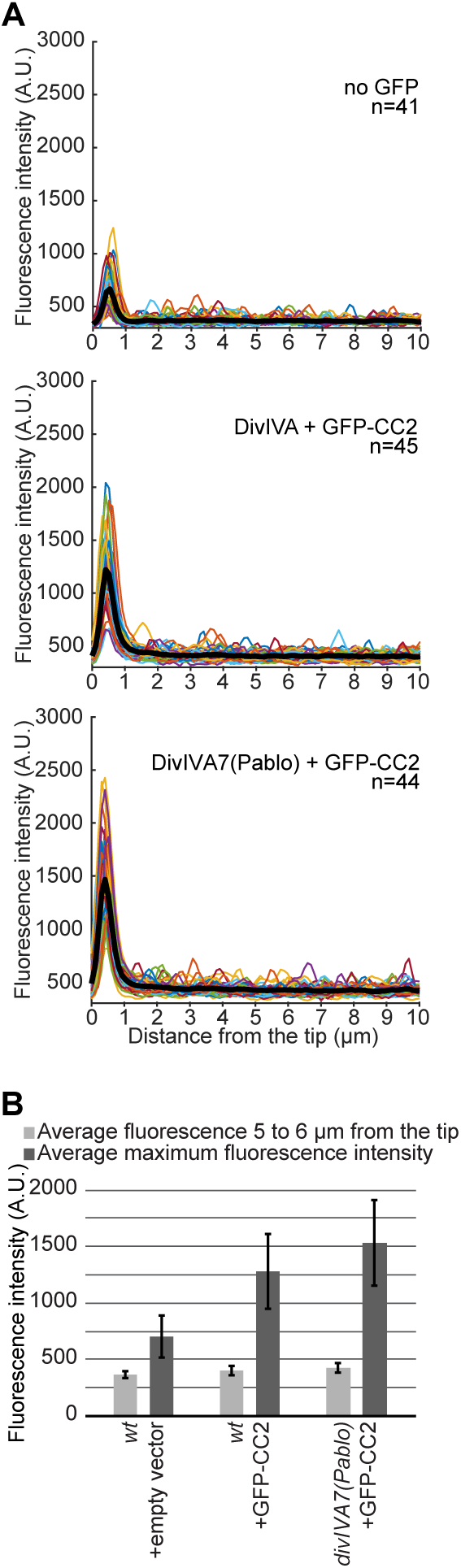
Fluorescence profiles of DivIVA(CC2)-GFP along apical regions of growing hyphae. **A.** Fluorescence profiles along individual hyphae (colored lines) and average fluorescence profiles (bold black line). The zero on the x-axis corresponds to the maximum intensity in the phase contrast image, in the halo just outside the hyphal tip. **B.** Average signal along the hyphae, 5-6 mm from the tip (light grey) compared to the maximum intensity (dark grey) displayed at the tip in the fluorescence profiles. Error bars show standard deviations.

It has previously been observed that conditions that cause growth arrest in *S. coelicolor* tend to trigger the disappearance of the DivIVA-based polarisome at the hyphal tips [11, 18, 22, 61]. The availability of the *divIVA7*(Pablo) mutant allowed us now to test whether phosphorylation of DivIVA is required for the disassembly or disappearance of polarisomes from the hyphal tips upon growth arrest. We tested specifically the bacitracin-induced growth arrest, since it is well established that AfsK-mediated DivIVA phosphorylation is strongly upregulated under such conditions (see Fig 2 and [22]). The wild-type parent and the *divIVA7*(Pablo) mutant, both expressing the DivIVA GFP-CC2 fusion protein, were grown in the microfluidic cell perfusion system as described above, and the response to addition of bacitracin was monitored. In both cases, extension of all observable hyphal tips arrested, and the apical fluorescence signal from the polarisome structure disappeared instantaneously (meaning it was gone at the same time point as arrest of hyphal tip extension was detected, using a 20 min interval between images)(S5 Fig). The response was indistinguishable between the two strains, showing that dismounting of polarisomes from stalled hyphal tips does not require DivIVA phosphorylation under bacitracin-induced growth arrest.

## Discussion

Streptomycetes are morphologically highly plastic and adapt their growth patterns and morphology to environmental conditions. The DivIVA-based polarisome and the activities at the hyphal tip are likely to be focal points for regulatory mechanisms that affect the morphological plasticity. We have previously shown that the Ser/Thr protein kinase AfsK is involved in such regulation and that it targets DivIVA in *S. coelicolor* [22]. In this paper, we identify a protein phosphatase, SppA, that dephosphorylates DivIVA, and we show that SppA is involved in the regulation of polar growth in *S. coelicolor*. The partial suppression of the microscopically observable phenotypes of the *sppA* mutant by *afsK* deletion suggests that that SppA and AfsK at least partially are in the same regulatory pathway.

SppA was previously reported to dephosphorylate serine, threonine and tyrosine residues in phosphorylated peptides *in vitro* [58], but none of its natural targets had been identified. Here, we provide evidence that SppA dephosphorylates DivIVA both *in vivo* and *in vitro*. Further, the strong effects of *sppA* deletion on tip extension rate, increased frequency of spontaneous growth arrests of individual tips, and overall effect on hyphal morphology and branching that we report here are likely the reasons for the slow growth of cultures of *sppA* mutants [58],

The mutant phenotype shows that *sppA* has non-redundant tasks that are not fulfilled by any of the other predicted protein phosphatases encoded by *S. coelicolor*. A 2004 survey revealed 55 genes coding for phosphatases predicted to target *O*-phosphorylated proteins in *S. coelicolor*: 4 predicted tyrosine phosphatases and 51 Ser/Thr phosphatases, out of which 2 (including SppA) are members of the phosphoprotein phosphatase (PPP) superfamily and 49 of the Mg^2+^- or Mn^2+^-dependent protein phosphatase (PPM) superfamily [62]. Of the 49 PPM-family genes, the major fraction are from one subfamily that appear to have expanded substantially in *Streptomyces* genomes (a similar number were found in *Streptomyces avermitilis*), with many genes found in the chromosomal arms that tend to contain genes for non-essential functions [63]. Nine of the PPM-family genes in *S. coelicolor* were in gene clusters together with an STPK gene, suggesting that they may be working together with a specific cognate kinase [62], and it was also speculated that many of the PPM-family-associated kinases may be involved in RNA polymerase sigma factor regulation [63]. For 30 of the 49 PPM-family proteins, the catalytic phosphatase domain is connected to additional domains of distinct predicted functions. Thus, even if there is a large expansion of the PPM-family proteins in the *Streptomyces* genomes, it is likely that many are dedicated to specific pathways or functions, and only have one or a few specific substrates. Out of the two PPP-family genes identified in *S. coelicolor* [62], *sco5973* is adjacent to a gene predicted to encode the RNA ribose 2’-O-methyltransferase Hen1 and was therefore suggested to encode a polynucleotide phosphatase involved in bacterial RNA repair [64]. Thus, SppA appears to be the only PPP-family protein phosphatase in *S. coelicolor*.

Typically, the number of protein phosphatase genes is lower than the number of kinase genes in both bacterial and eukaryotic genomes. A typical example in bacteria is *M. tuberculosis*, which encodes 11 STPKs and one single STP [57]. Importantly though, this low number of phosphatases does not mean that they are promiscuous enzymes that dephosphorylate a host of different substrate proteins. Mammalian phosphatases of the PPP family, such as the most abundant PP1- and PP2A-type, are catalytic subunits in heteromeric enzymes with additional subunits that control activity and confer substrate specificity, including cell-type or compartment specificity [65–67]. The streptomycete SppA has no additional domain that could confer substrate selectivity or control its activity, but it may have interaction partners fulfilling that role. Thus, despite the subcellular distribution that we found of SppA throughout the hyphae, its activity or specificity could locally be directed by other proteins.

Our previous work showed that AfsK phosphorylates DivIVA in *S. coelicolor*, and that no DivIVA phosphorylation is detectable in an *afsK* mutant [22]. Based on the identification of one threonine and four serine residues that are phosphorylated in DivIVA [23], a phospho-ablative mutant *divIVA* allele that abolishes detectable phosphorylation was constructed, and allowed for example the demonstration that delocalization of DivIVA from hyphal tips upon bacitracin-imposed growth arrest is unaffected by DivIVA phosphorylation. Interestingly, this *divIVA7*(Pablo) mutant yielded a similar effect on hyphal branching during vegetative growth as previously reported for the *afsK* mutant, with longer average distance from hyphal tips to sites where first lateral branches emerge (tip-to-branch distance). Thus, it can now be concluded that this phenotypic effect of *afsK* is mediated by phosphorylation of the polarisome protein DivIVA.

On the other hand, the phenotypic effects of the *sppA* deletion were not suppressed by the *divIVA7*(Pablo) allele, showing that although there are elevated levels of DivIVA phosphorylation in the *sppA* mutant compared to the wild type, these direct effects on DivIVA are not causing the slow tip extension and altered branching seen in the mutant. Importantly though, *afsK* deletion partially suppresses the *sppA* phenotype, suggesting that AfsK has additional substrates that are dephosphorylated by SppA and that when hyper-phosphorylated lead to effects on apical growth and branching. Apart from DivIVA, two other targets of AfsK have previously been reported. The transcriptional regulator AfsR is involved in control of antibiotic production and is phosphorylated by AfsK in *in vitro* assays [55, 68, 69]. Further, the replication initiator protein DnaA is phosphorylated in actively growing mycelium of *S. coelicolor*, and this phosphorylation is dependent on *afsK* [70]. A phospho-mimic mutant form of DnaA showed elevated ATPase activity and decreased affinity for the *oriC* region in *in vitro* binding assays [70]. However, there is no known connection between DnaA and hyphal tip extension and cell shape, making it unlikely that DnaA would mediate the *afsK*-dependent effect of *sppA* on hyphal growth. Instead, we suggest that AfsK and SppA share other target proteins and that these would be highly interesting to identify with respect to control of polar growth and cell shape determination. In addition, based on the incomplete suppression by *afsK*, SppA is likely to have additional targets that affect growth but are phosphorylated by other kinases than AfsK.

Overall, our findings emphasize the importance of Ser/Thr phosphorylation for control of growth and cell cycle-related processes in *Streptomyces* and Actinobacteria. The pathway that we are starting to uncover in *Streptomyces* is unusual in that SppA is a PPP-family phosphatase and that AfsK has a sensory domain consisting of PQQ repeats, predicted to form a β-propeller structure [46]. In most previous reports about cell wall and growth-related regulation in other Gram-positives, the phosphatases are from the PPM family and the kinases contain peptidoglycan- or lipid II-binding PASTA domains [25-29, 40, 56, 57]. AfsK activity is stimulated when peptidoglycan synthesis is arrested by bacitracin or vancomycin [22], and it will be highly interesting to determine exactly what is sensed by AfsK and how this leads to activation of kinase activity. Further, the role of the phosphatase SppA in hyphal growth and morphogenesis and its substrate specificity should be further investigated.

## Materials and Methods

### Strains, plasmids and media

All bacterial strains used in this study are listed in Table 1. Plasmids and oligonucleotides are listed in S1 and S2 Tables, respectively. *E. coli* strains were grown as appropriate in Lysogeny broth (LB) or on LB agar [71] supplemented with kanamycin (50 µg/ml) or apramycin (50 µg/ml), or in LB without salt supplemented with hygromycin (50 µg/ml). Unless specified otherwise, *S. coelicolor* strains were grown as appropriate on Soya Flour Mannitol (SFM) agar [72] supplemented with apramycin (25 µg/ml), hygromycin (25 µg/ml), viomycin (30 µg/ml), or in TSBS (BD Bacto Tryptic Soy Broth, supplemented with 17% sucrose) in flasks with stainless steel spirals. Transfer of plasmids by conjugation from *E. coli* to *S. coelicolor* was done as described previously [72], using *E. coli* strain ET12567/pUZ8002 as conjugation donor.

When inoculation of cultures with a precise number of *S. coelicolor* CFUs was needed, the spores were harvested from 6- to 8-days-old bacterial lawn on SFM plates, filtered and concentrated in 20% glycerol as previously described [72], then aliquoted by 50 µl and frozen at −80°C. One aliquot was then used to determine the spore titer: serial dilutions of the aliquot were made in 0.005% Tween 80 to avoid clumping of the hydrophobic spores, and plated on SFM. Cultures for investigation of mycelial morphology were grown on cellophane membranes placed on the surface of agar medium, as described previously [61], before the cellophanes were removed and mounted on agarose-coated slides for microscopical investigation.

### Purification of His_6_-SppA

*E. coli* strain TUNER(DE3)/pKF461 was cultivated in 1 L of LB medium, 37°C to OD_600_ of 1, when isopropyl-β-D-thiogalactoside (IPTG) was added to 0.33 mM and the culture thereafter incubated at 16°C for 20 hours. Harvested cells were resuspended in 25 ml of 50 mM sodium phosphate buffer, pH 8.0, 300 mM NaCl, 20 mM imidazole, supplemented with EDTA-free Complete Protease Inhibitor cocktail (Roche) and passed three times through a French pressure cell at 18,000 psi. The resulting lysate was centrifuged at 200,000 x g, 60 min, 4°C, and the supernatant was applied on a 1 ml HisTrap HP column (GE Healthcare) equilibrated with the same buffer. The column was washed with the same buffer until a stable UV absorption signal was observed. The bound protein was eluted with a gradient of 20-500 mM imidazole in 50 mM sodium phosphate buffer, pH 8.0, 300 mM NaCl over 20 column volumes, followed by 5 column volumes of 50 mM sodium phosphate buffer, pH 8.0, 300 mM NaCl, 500 mM imidazole. Fractions were collected and those that contained most SppA, as seen by SDS-PAGE analysis and pNPP assays (see below), were pooled and further purified by gel filtration on a HiLoad 26/600 Superdex 75 pg column. Fractions with highest phosphatase activity were pooled, mixed with glycerol to a final concentration of 10% and stored in aliquots at −80°C. Assays of phosphatase activity using para-nitrophenylphosphate (pNPP) substrate showed that the protein did not lose activity during storage under these conditions. Phosphatase assays were carried out at 30°C in 50 mM Tris-HCl, pH 8.0, 2 mM MnCl_2_, and 10 mM pNPP, and detecting absorbance at 405 nm, as described previously [58].

### Dephosphorylation of DivIVA *in vitro*

Phosphorylated DivIVA (phospho-DivIVA) was obtained by co-producing the kinase domain of AfsK with DivIVA, fused to an intein and a chitin-binding domain, in *E. coli*. *E. coli* strain ER2566 carrying plasmids pKF135 and pKF358 was cultivated in 1 L of LB medium to an OD_600_ of 0.5, whereafter IPTG was added to a final concentration of 0.3 mM and the culture was incubated 18 h at 20°C. Cells were harvested by centrifugation and resuspended in 25 ml of 20 mM HEPES, pH 8.0, 500 mM NaCl, 1 mM EDTA, supplemented with EDTA-free Complete Protease inhibitor cocktail (Roche), and passed twice through a French pressure cell at 18,000 psi. The resulting lysate was centrifuged at 200,000 x g, 60 min, 4°C and the supernatant applied on a 6 ml gravity flow column with Chitin resin (New England Biolabs), equilibrated with 20 mM HEPES, pH 8.0, 500 mM NaCl, 1 mM EDTA. The column was then washed with 10 column volumes of 20 mM HEPES, pH 8.0, 500 mM NaCl, 1 mM EDTA, 0.1% (w/v) of Triton X-100, followed by 10 column volumes of 20 mM HEPES, pH 8.0, 1 M NaCl, 1 mM EDTA, 1% (w/v) of Triton X-100, and thereafter 3 times 2 column volumes of 20 mM HEPES, pH 8.0, 500 mM NaCl, 1 mM EDTA, 5 mM ATP, 10 mM MgCl_2_ with 3 min incubation after each 2 volumes. Finally, the column was washed with 4 column volumes of 20 mM HEPES, pH 8.0, 500 mM NaCl, 1 mM EDTA, followed by 20 ml of freshly prepared cleavage buffer (20 mM HEPES, pH 8.0, 500 mM NaCl, 1 mM EDTA, 50 mM dithiothreitol). The column was then plugged and incubated overnight at 4°C, whereafter the protein was eluted with 3 column volumes of 25 mM Tris-HCl, pH 7.5, 150 mM NaCl, 10% glycerol. Fractions containing pure phospho-DivIVA, as determined by SDS-PAGE, were pooled, dialysed against the same buffer, and stored in aliquots at −80°C.

Dephosphorylation of phospho-DivIVA by SppA was monitored under the following conditions: 50 mM Tris-HCl, pH 8.0, 2 mM MnCl_2_, 15 mM NaCl, 0.21 μM phospho-DivIVA, incubated at 55°C. Reactions were started by adding His_6_-SppA to a final concentration of 35 nM. Control reactions with addition of 20 mM EDTA or without SppA were also included. Reactions were stopped at different time points by addition of SDS loading buffer and heating to 100°C, 5 min. Proteins were separated by SDS-PAGE and proteins detected by Western blotting with anti-DivIVA antiserum as described previously [73]. The reaction conditions had first been optimised with respect to pH, NaCl concentration, and temperature, with results consistent with a previous report on the phosphatase activity of SppA [58].

### Allelic replacement of *divIVA*

The allelic replacement of *divIVA* by a phospho-ablative allele with five codons (for residues T304, S309, S338, S344, and S355) changed to alanine codons, *divIVA7*(Pablo), was obtained using a *codA* counter selection system previously described by Dubeau *et al*. [74]. The mutated *divIVA7*(Pablo) allele, flanked by 2 kb of genomic sequence upstream and downstream was cloned into the *codA*-containing vector pMG303M, giving pKF474 (see plasmid table for construction details). This plasmid was then introduced in *S. coelicolor* strain M600 by protoplast transformation [72]. Transformants with integration of the mutated plasmid by homologous recombination were obtained by selection with hygromycin. The isolated transformants were grown to sporulation on non-selective SFM plates to allow for excision of the plasmid by homologous recombination. Serial dilutions of spores were then plated on counter-selective medium (minimal medium supplemented with 50 µg/ml 5-fluorocytosine, 5FC) to allow growth only of strains that had lost the integrated plasmid, as described previously [74]. The *codA*-carrying clones convert 5FC to 5-fluoro-uracil (5FU) and die from it. At high cell density, this releases enough 5FU in the plates to also kill the clones without *codA*; candidate excision clones were thus picked on the lower density plates. They were then streaked on SFM to allow sporulation and from there clean-streaked on SFM to segregate genetically homogenous clones. These colonies were tested for resistance to 5FC and sensitivity to hygromycin (verifying loss of plasmid). The genomic DNA of the positive candidates was then extracted (see below) and the region containing the target mutations was PCR-amplified and sequenced. One strain carrying the *divIVA7*(Pablo) allele was named K365.

### Extraction of genomic DNA from *S. coelicolor*

All centrifugation steps were done at maximum speed in a benchtop centrifuge. Mycelia were pelleted from 2 ml of overnight liquid culture. 200 µl of Zirconia/silica beads (BioSpec products) were added to the pellet, as well as 100 µl lysis buffer (2% Triton, 1% SDS, 100 mM NaCl, 10 mM Tris-HCl, pH 8, 1 mM EDTA) and 100 µl Phenol:Chisam 1:1 (Chisam, Chloroform:Isoamyl alcohol 24:1). The cells were broken with a FastPrep-24 bead-beater (MP Biomedicals) by 4 to 6 cycles of 30 sec at 4 m/s. 200 µl 1X SSC (150 mM NaCl, 15 mM Trisodium citrate, pH 7) was added and mixed into the tube, then the sample was centrifuged for 3 min. The aqueous phase was harvested and completed to 600 µl with TE buffer (10 mM Tris-HCl, pH 7.5, 1 mM EDTA). 100 µl of 5 M NaCl was added and the sample mixed thoroughly before adding 80 µl of CTAB-NaCl (10% hexadecyltrimethyl ammonium bromide, 0.7 M NaCl) pre-warmed at 65°C. The samples were incubated at 65°C for at least 10 min, then an equal volume of Chisam was added and the samples mixed thoroughly before being incubated on the bench for a few minutes. After centrifugation for 10 min, the aqueous phase was harvested and mixed with 0.6 volumes of isopropanol, then centrifuged for 15 min. The supernatant was discarded and the pellet washed with 500 µl of 70% Ethanol at −20°C. The pellet was then dried and finally dissolved in 50 µl TE supplemented with 100 µg/ml RNase A and incubated at 37°C for 30 min.

### Time-lapse microscopy in microfluidic growth chambers

The method is adapted from Schlimpert *et al*. [59]. Briefly, a CellASIC B04A-03 microfluidic plate was rinsed and primed with the appropriate media, typically TES:DGM (one part 0.05 M TES buffer pH 8, one part Double Strength Germination medium [72]) in wells 1, 2, 6 and 8, TSB in wells 3, 4 and 5. *S. coelicolor* spore aliquots were diluted in TES:DGM down to 5×10^6^ CFU/ml and 50 µl of this suspension was loaded into the cell inlet well (well 8). If necessary, the TSB used for priming in wells 3 to 5 was replaced with TSB supplemented by active compounds (e.g. thiostrepton at 1 μg/ml or bacitracin at 50 μg/ml), as appropriate. The spores were then loaded from the inlet well 8 to the growth chamber with the following program: V8, 2 psi, 15 sec / V6+V8, 3 psi, 20 sec / V6, 1 psi, 30 sec. The spore density in the chamber was controlled under the microscope, with an aim of 1 to 3 spores at depth 0.9 µm per field of 200×200 µm. Typical microfluidic program is: 2 psi, 8h, TES:DGM / 2 psi, 15h, TSB. The microscope incubator was set at 30°C. The spores were allowed to germinate overnight (13 to 14h) before starting the observation. The observation was done with an inverted Zeiss Axio Observer.Z1 controlled through ZEN software, using a 63X phase contrast objective and Zeiss filter sets 38HE (GFP) or 63HE (mCherry), as appropriate. Zeiss Definite Focus was used to correct Z-drift every 10 sec between the time points, with a reference position in one of the medium channels. Time interval was typically 15 min.

### Microscopy in agarose growth chambers

The appropriate growth medium was supplemented with 1% agarose (electrophoresis grade), melted and casted in a custom-made support ([11], design available upon request), creating an agarose pad held in a frame. To get the mycelia to grow as flat as possible, a cellophane piece was layered on top of the agar pad, then a suspension of spores or young mycelia was inoculated onto the cellophane and covered with a coverslip. To allow for proper oxygenation of the medium while limiting desiccation, the back of the agar pad is covered with a gas-permeable membrane, as described previously [11]. The assembled growth chamber was incubated in a humidified box at 30°C until observation.

### Image analysis

Image analysis was conducted mainly using Fiji [75]. Prior to any analysis, time-lapse series were re-aligned to correct for XY jitter with the “Correct 3D drift” plugin [76].

Tip extension rates were obtained from manual tracking of the tips with the TrackMate plugin [77] from their emergence until they are lost out of the field, overlapped by other hyphae, or until they turn too sharply for a straight line to represent their elongation. The age of the hyphae (time relative to the start of the track) at each time-point was then calculated using Matlab R2016B (Mathworks) and the distribution of extension rates for hyphae 1 to 3h old was calculated using Microsoft Excel. Statistical analysis was carried out using PAST [78].

The distance from a tip to a proximal new branch was obtained by manually tracing the hyphae with a broken line from the base of the new branch to the tip of the hypha at the time point when the new branch was fully mature and started extending rapidly (1 to 4 µm long). The length of the line was then measured and exported to MS Excel for calculation of the distribution. Statistical analyses were done using PAST [78].

Fluorescence profiles from the tips of hyphae were measured with the help of a custom ImageJ macro, then processed in Matlab using a custom script to align them according to the maximum intensity in the phase contrast channel, which would be just a little outside the tip.

### Detection of DivIVA phosphorylation by Western blot

The electrophoretic mobility shift of DivIVA due to its phosphorylation was detected by Western blot as previously described [22], with only minor modifications. Briefly, mycelia from 10 to 20 ml of culture were harvested by centrifugation (10 min, 3.000 x g), rinsed twice in lysate buffer and resuspended in lysate buffer (twice the volume of the cell pellet). They were then broken by bead-beating, and the clear lysate was obtained by centrifugation. The total protein concentration of the clear lysate was quantified using BioRad DC protein assay. 10 µg of total proteins were separated on a BioRad precast stain-free 12% acrylamide SDS-PAGE. The StainFree gel was activated 2.5 min and imaged with a BioRad ChemiDoc MP, then transferred to a PVDF membrane. The membrane was blocked with 5% powder milk in PBS-Tween, then incubated with 1:5,000 diluted anti-DivIVAsc anti-serum from rabbit [73], washed three times in blocking solution, incubated with pig anti-rabbit IgG cross-linked to horseradish peroxidase (1:1,000; DakoCytomation), and washed three times in PBS-Tween. The blot was incubated with SuperSignal West Pico chemiluminescence substrate (Thermo scientific) and imaged with a BioRad ChemiDoc MP for both chemiluminescent (DivIVA) and StainFree (total proteins) signals.

## Acknowledgements

The Lund Protein Production Platform (LP3) is acknowledged for assistance with protein production and purification. We thank Elisabeth Barane for excellent technical assistance, and Juri Kazakevych, Antje Hempel, Fan Yang, Algirdas Miksys, Malou Restedt, and Natalia Berges for contributions to strain constructions and preliminary experiments. We are grateful to Paul Dyson, Ryszard Brzezinski, Matt Bush, and Mark Buttner for gifts of plasmids and strains. Erik Svensson and Jörgen Ripa are acknowledged for advice on statistical analysis and Matlab, respectively.

## Supporting information

S1 Table. Plasmids used in this study

S2 Table. Oligonucleotide primers used in this study

S1 Fig. *In vivo* dephosphorylation of DivIVA

S2 Fig. Subcellular SppA localisation

S3 Fig. Mycelial morphology in microfluidic cell perfusion system

S4 Fig. No effect of *divIVA7*(Pablo) on colony phenotype on different media

S5 Fig. Phosphorylation of DivIVA is not required for dissipation of apical polarisomes upon or recovery from bacitracin-induced growth arrest

S1 Movie. Mycelial growth of wild type strain M600 in microfluidic cell perfusion system.

S2 Movie. Mycelial growth of *sppA* mutant strain K350 in microfluidic cell perfusion system.

